# Adeno-associated virus delivered Cre recombinase as a versatile strategy for modeling focal neuronal lesions

**DOI:** 10.1101/2025.05.20.655018

**Authors:** Margaux Giraudet, Emma Cloarec, Frédéric Gambino, Elisabete Augusto

**Affiliations:** Institut Interdisciplinaire de Neurosciences (IINS), Univ. Bordeaux, CNRS, IINS, UMR 5297, F-33000 Bordeaux, France; Centre Broca Nouvelle-Aquitaine, 146, rue Léo-Saignat, 33076 Bordeaux, France

## Abstract

Lesion studies remain central to neuroscience, offering key insights into brain-behavior relationships. Excitotoxic agents, though widely used, often produce inconsistent, non-specific, and ethically complex outcomes. In contrast, optogenetics and chemogenetics enable precise, reversible manipulations but pose technical and interpretational challenges, including cost and non-physiological activation. There remains a need for robust, controlled, and ethical lesion methods. Here, we present Cre recombinase expression as a reproducible alternative to excitotoxic lesions in mice. Using high titration adeno-associated viral (AAV) vectors, we show that Cre expression induces focal neurodegeneration across brain regions, affecting both excitatory and inhibitory neurons. This effect is independent of viral serotype or fluorescent tags and is not visible with standard staining but is revealed by neuronal markers. The resulting neuronal loss is followed by glial remodeling and significant behavioral alterations. AAV-mediated Cre expression thus provides a powerful, genetically targeted lesion model that bridges classical and modern approaches to circuit manipulation.

## INTRODUCTION

Lesion studies have long served as a foundational approach in neuroscience, providing crucial insights into brain-behavior relationships^1,2^. By investigating the effects of permanent damage to, or removal of, specific brain regions, these approaches have been instrumental in identifying the neural substrates of diverse behaviors^3^ and in developing models of neurodegenerative disorders^4–6^. For example, the use of 6-hydroxydopamine (6-OHDA), a neurotoxin that selectively targets and induces degeneration of dopaminergic neurons, has become a widely used and invaluable tool for modeling Parkinson’s disease in animal studies^5^.

In recent years, however, the emergence of optogenetic and chemogenetic techniques has transformed our capacity to causally link neural activity within defined circuits to specific behavioral outcomes. These modern approaches offer temporally precise, reversible manipulation of neuronal activity with remarkable cell-type and circuit specificity^7^. However, these techniques also introduce a range of technical and interpretational challenges. Optogenetic for instance requires the implantation of light delivery systems, which can be more technically demanding and costly than traditional lesion methods. Artificial optogenetic stimulation may also evoke non-physiological firing patterns that do not accurately reflect natural neural activity^8–10^, and can trigger rebound excitation when stimulation ends^11–13^, potentially resulting in artifactual behavioral effects. These issues are further complicated by the limited penetration of visible light through scattering brain tissue^14^. As such, classical lesion approaches continue to offer important strategic advantages, particularly in the early stages of investigation, by providing a fast, cost-effective, and robust method for probing brain–behavior relationships.

Classic approaches for producing chronic lesions involve the use of excitotoxic agents such as kainic acid, quinolinic acid, ibotenic acid, or glutamate^1,2^. These compounds overstimulate glutamate receptors, leading to localized neuronal death at the injection site while typically sparing passing axonal fibers. Despite their utility, classical excitotoxic agents often results in inconsistent and unpredictable lesion outcomes across different brain areas, complicating interpretation and reproducibility^1,2,15^. Furthermore, these agents lack cell-type and circuit specificity, often affecting diverse neuronal populations and thus confounding causal inferences. More critically, excitotoxic lesions are inherently invasive and are frequently associated with severe side effects, including seizures and poor post-operative recovery, which can be lifethreatening. These risks raise significant ethical concerns due to the potential for substantial distress or long-term harm to the animal. Collectively, these limitations highlight the need for an alternative method to induce lesions in the CNS that is more controlled, reproducible, and ethically sound.

Recent studies have proposed the use of Cre recombinase as a potent alternative to excitotoxic agents for inducing localized brain lesions^16–18^. Cre recombinase has revolutionized genetic research by enabling tissue- and cell-type-specific gene manipulation. It operates by introducing *loxP* sites flanking the gene of interest and then expressing the Cre enzyme, which recognizes and excises the DNA between these sites^19^. However, Cre recombinase can inadvertently recognize pseudo-loxP-like sequences in the genome, leading to unintended DNA damage and, in some cases, programmed cell death^20^. Interestingly, this cytotoxic effect has been observed serendipitously in dopaminergic neurons^16,17^ and thalamic neurons^18^, suggesting its potential as a tool for targeted neuronal ablation.

Here, we demonstrate that adeno-associated virus (AAV)-mediated expression of Cre recombinase induces severe neurodegeneration at the injection site, an effect that occurs independently of the associated serotype or fluorescent tag. Our findings indicate that high levels of Cre expression are the primary driver of this neurodegeneration, which was not readily detectable using standard DAPI staining, due to glial cell infiltration masking neuronal loss. Instead, the neurodegeneration was clearly revealed through immunostaining with neuronal markers. Notably, we demonstrate that elevated Cre expression induces focal lesions across multiple brain regions, impacting both excitatory and inhibitory neuronal populations. These lesions are followed by glial-mediated tissue remodeling and result in pronounced behavioral alterations. Together, these results highlight the utility of AAV1-mediated Cre expression as a promising tool for inducing chronic, localized, and neuron-specific lesions within the central nervous system.

## RESULTS

### Injection of AAV1-CRE induces a rapid and robust Cre recombinase activity, followed by a localized and progressive loss of neurons

We first tested whether injection of an adeno-associated virus expressing Cre recombinase (AAV1-CRE) induces neuronal loss (**Fig.1**). AAV1-CRE was delivered unilaterally into the medio-dorsal thalamic nucleus (MD) (**Fig.1A**). We quantified neuronal survival at multiple time points post-injection by performing post-mortem immunohistochemistry for NeuN, a widely used marker of mature neurons in the forebrain^21,22^. We used the DeepSlice software^23^ and the Quint workflow^24^ to register optical sections and delineate precise anatomical subdivisions of the MD based on the Allen Brain Institute taxonomy. This enabled accurate and automated quantification of NeuN-positive cells specifically within the injected MD (**Fig.1A, B**). We found no significant difference in the number of NeuN+ cells between hemispheres in sham controls (mean ± standard deviation (sd); lesion: 2520 ± 324.1; non-lesion: 2525 ± 214.9, p=0.9546, paired t-test). Therefore, we normalized the number of NeuN+ cells in the injected MD to those in the contralateral, noninjected MD, which served as an internal control for signal normalization (**Fig.1A, methods**).

**Figure 1.**
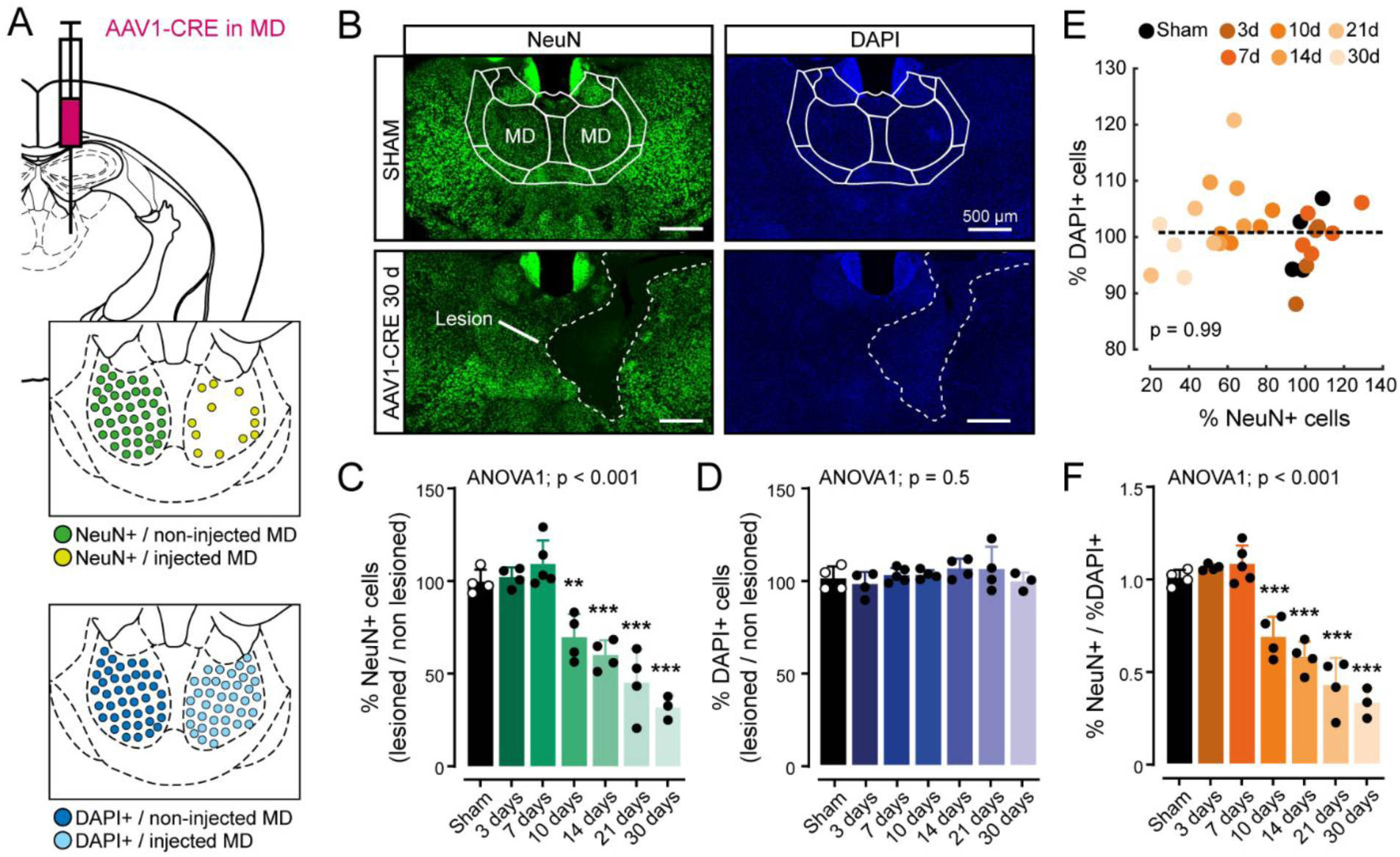
Cre-mediated progressive neuronal loss. **A)** Top: schematic representation of the stereotaxic delivery of AAV1-CRE into the right MD. Bottom: schematic representation of the results presented in C-D. **B)** Top: representative epi-fluorescent image of a coronal brain slice aligned to the Allen brain atlas taken from a mouse not injected with AAV1-CRE into the MD (sham). Bottom: representative epi-fluorescent image of a coronal brain slice taken from a mouse injected with AAV1-CRE into the right MD 30 days before sacrifice (30 d). Green: NeuN, blue: DAPI. **C)** % of NeuN-positive cells in the injected side and normalized by the number of NeuN-positive cells in the non-injected side at different time points after AAV1-CRE injection into the right MD. Error bars are sd. **D)** % of DAPI-positive cells in the injected side and normalized by the number of DAPI-positive cells in the non-injected side at different time points after AAV1-CRE injection into the right MD. Error bars are sd. **E)** Correlation between % of NeuN-positive cells at the different time points post-injection represented in C, and the % of DAPI-positive cells represented in D. Each dot corresponds to a mouse. Dashed line is the best fit using a simple linear regression. **F)** % of NeuN-positive cells in the injected side and normalized by the number of DAPI-positive cells in the same region at different time points after AAV1-CRE injection into the right MD.

At 3- and 7-days post-injection, the fraction of NeuN-positive cells in the injected MD did not differ from that of sham-injected mice. However, by 10 days post-injection, NeuN-positive cell counts were significantly reduced by an average of 30 % compared to sham mice. This neuronal loss progressed significantly over time, with average reductions of 40% at 14 days, 55% at 21 days, and 68% at 30 days post-injection (**Fig.1C**; one-way ANOVA followed by Dunnett’s multiple comparison test, p<0.0001). In contrast, the total number of DAPI-positive cells remained unchanged across all evaluated time points (**Fig.1B, D**; one-way ANOVA followed by Dunnett’s multiple comparison test, p=0.5076). Accordingly, no correlation was observed between NeuN-positive cells and DAPI-positive cells at the injected site (**Fig.1E**; Pearson’s correlation, p = 0.9867, r^2^ < 0.00001). Consequently, the percentage of NeuN-positive cells normalized to DAPI (**Fig.1F**) mirrored the trend observed in absolute NeuN percentages (**Fig.1C**), revealing a progressive and specific decline in neuronal populations over time (mean fraction of NeuN+ positive cells ± sd; Sham; 1.002 ± 0.043; AAV1-CRE; + 3 days: 1.06 ± 0.017; +7 days: 1.08 ± 0.1; + 10 days: 0.68 ± 0.11; 14 days: 0.57 ± 0.08; +21 days: 0.42 ± 0.15; + 30 days: 0.33 ± 0.08). These results indicate that DAPI staining does not reliably reflect neurodegeneration, as the loss of NeuN-positive neurons is paralleled by a proportional increase in non-neuronal cell types within the lesioned brain tissue.

We next characterized the temporal dynamics of Cre expression, by quantifying the percentage of Cre-positive cells using immunohistochemistry against Cre (**Fig.2A, B**). Although neurodegeneration became apparent around 10 days post-injection (see **Fig.1**), Cre recombinase expression was detectable as early as 3 days post-injection, with a marked and significant increase in the proportion of Cre-positive cells compared to sham controls (Sham: 0.0025 ± 0.005; AAV1-CRE: 0.42 ± 0.32). The expression of Cre continued to rise at 7 days (0.53 ± 0.25) and peaked at 10 days post-injection (0.64 ± 0.29). Beyond this point, however, the percentage of Cre-positive cells gradually declined (14 days: 0.28 ± 0.12; 21 days: 0.13 ± 0.07; 30 days: 0.18 ± 0.02) (**Fig.2C**). Consequently, the progressive decline in Cre-positive cells showed a significant correlation with neuronal loss (**Fig.2D**, Pearson’s correlation, p = 0.03, r^2^=0.2026), indicating that Cre-expressing neurons underwent selective degeneration over time, thereby supporting the notion that Cre recombinase exerts neurotoxic effects.

**Figure 2.**
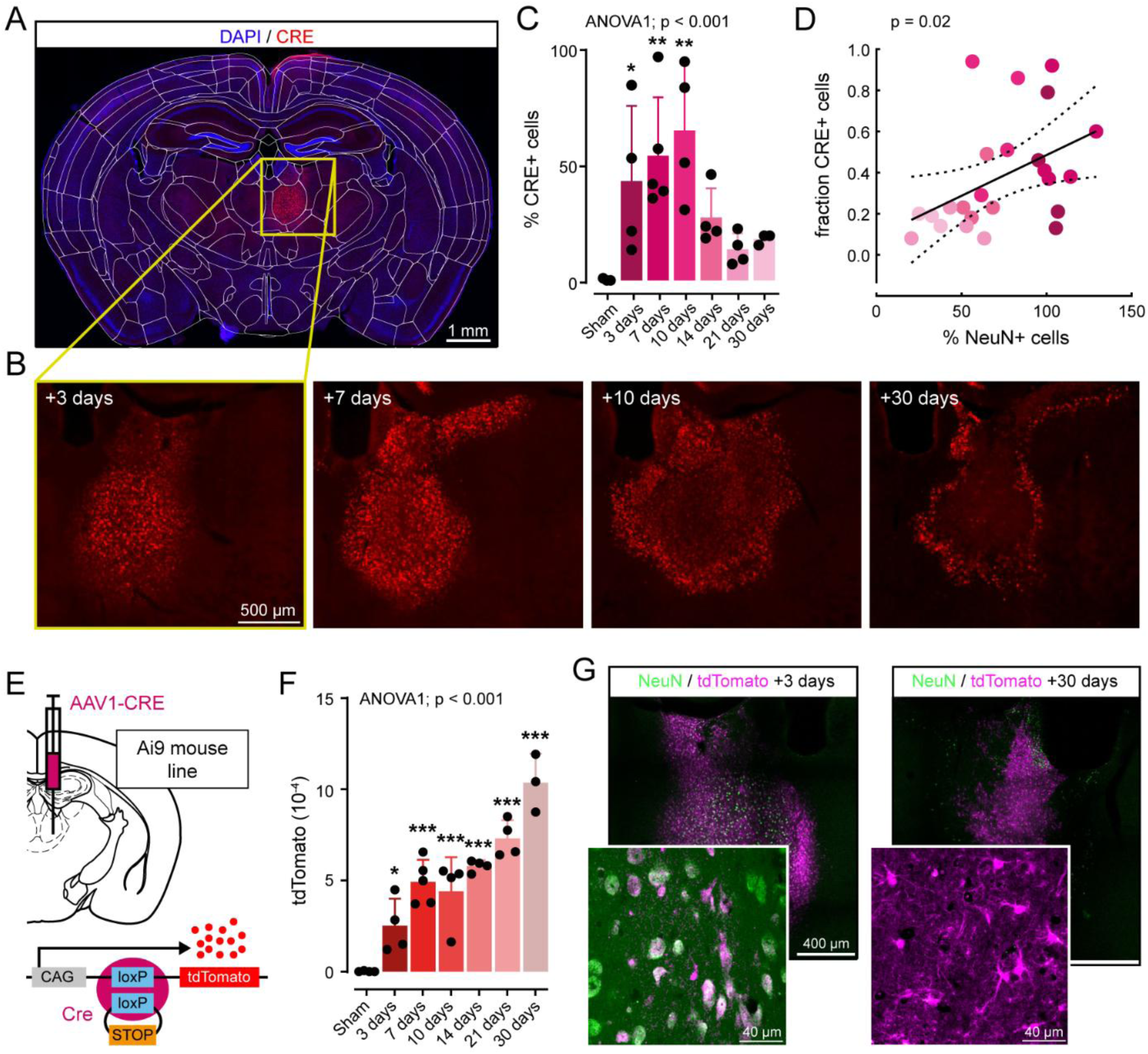
Cre-mediated Cre recombinase activity. **A)** Representative epi-fluorescent image of a coronal brain slice aligned to the Allen brain atlas taken from a mouse injected with AAV1-CRE into the MD of the right hemisphere, 3 days before sacrificed. Blue: DAPI, red: Cre. **B)** Zoomed image of the MD region represented by the yellow square in A), injected at different time points with AAV1-CRE in the MD of the right hemisphere. **C)** % of Cre-positive cells (normalized by DAPI-positive cells) in the injected side of the MD at different time points post-injection. Each dot corresponds to a mouse and error bars are sd. **D)** Correlation between % of NeuN-positive cells (normalized to NeuN -positive cells in the non-lesion side, as in figure 1C) in the MD at the different time points post-injection and the Cre-positive cells (normalized to DAPI-positive cells in the non-lesion side) in the MD. Each dot corresponds to a mouse represented in C). Filled line is the best fit using a simple linear regression. **E)** Schematic representation of the stereotaxic delivery of AAV1-CRE into the right MD of Ai9Tomato mice. **F)** Quantification of tdTomato fluorescence in the right MD at different time points of AAV1-CRE post-injection. **G)** Left: Zoomed images with different magnifications of the right MD 3 days post-injection of AAV1-CRE in the MD of the right hemisphere. Right: Zoomed images with different magnifications of the right MD 30 days post-injection of AAV1-CRE in the MD of the right hemisphere. Green: NeuN, purple: tdTomato.

To determine whether Cre recombinase could effectively mediate off-target recombination, and thereby induce neuronal death, prior to this decline, we injected AAV1-CRE into Ai9 TdTomato reporter mice (**Fig.2E**), which express TdTomato following Cre-mediated excision of a loxP-flanked stop cassette (**Fig.2E**). We quantified TdTomato fluorescence as a readout of recombination efficacy, which was significantly increased in AAV1-CRE-injected mice as early as 3 days post-injection compared to sham controls, where no TdTomato signal was putatively detected under the same imaging conditions (mean ± sd; Sham: 191.5 ± 322.5, AAV1-CRE 3 days: 25148 ±14971; p=0.04, one-way ANOVA followed by Dunnett’s multiple comparison test) (**Fig.2F**). These findings confirm the rapid expression of fully functional Cre recombinase following viral injection. Intriguingly, TdTomato signal continued to increase significantly at each time point measured (**Fig.2F**), even though the number of Cre+ cells declined after 10 days (**Fig.2A-C**). This suggests ongoing tissue remodeling, with other cell types beginning to display TdTomato and replacing the degenerated neurons. Consistent with this, while all TdTomato+ cells at 3 days post-injection exhibited neuronal morphology (**Fig.2G**, left), by 30 days, TdTomato was predominantly observed in cells displaying clear glial morphology **(Fig.2G**, right).

### Injection of AAV1-CRE induces neuronal loss followed by glial-mediated tissue remodeling across multiple brain regions

Dying neurons typically stimulate the activation and proliferation of glial cells at the lesion site, resulting in tissue remodeling and the formation of a persistent glial scar^25^. Here, we tested whether Cre-induced neuronal loss induced the formation of a glial scar by characterizing the temporal dynamic of GFAP expression (**Fig.3A**), a widely used marker for astrocytes^26,27^. We quantified the number of GFAP+ cells in the MD at various time points after the injection (**Fig.3B**), along with NeuN+ and DAPI+ cells as described above (see **Fig.1**). Compared to sham-injected mice, in which no GFAP signal was detected, the fraction of GFAP+ cells increased over time starting between 10- and 14-days post-injection (Sham: 6.5 ± 1.7; AAV1-CRE: 3 days: 7.1 ± 7.4, 7 days: 6.0 ± 4.0, 10 days: 4.8 ± 4.3, 14 days: 25.0 ± 15.4, 21 days: 66.5 ± 30.8, 30 days: 89.7 ± 9.0; p <0.0001) (**Fig.3C**). This increase indicates astrocyte proliferation at the injection site, which occurs after the onset of neurodegeneration around day 10. Accordingly, we observed a significant correlation between the decrease in NeuN+ cells and the increase in GFAP+ cells (**Fig.3D**, Pearson’s correlation, p < 0.0001, r^2^=0.656).

**Figure 3.**
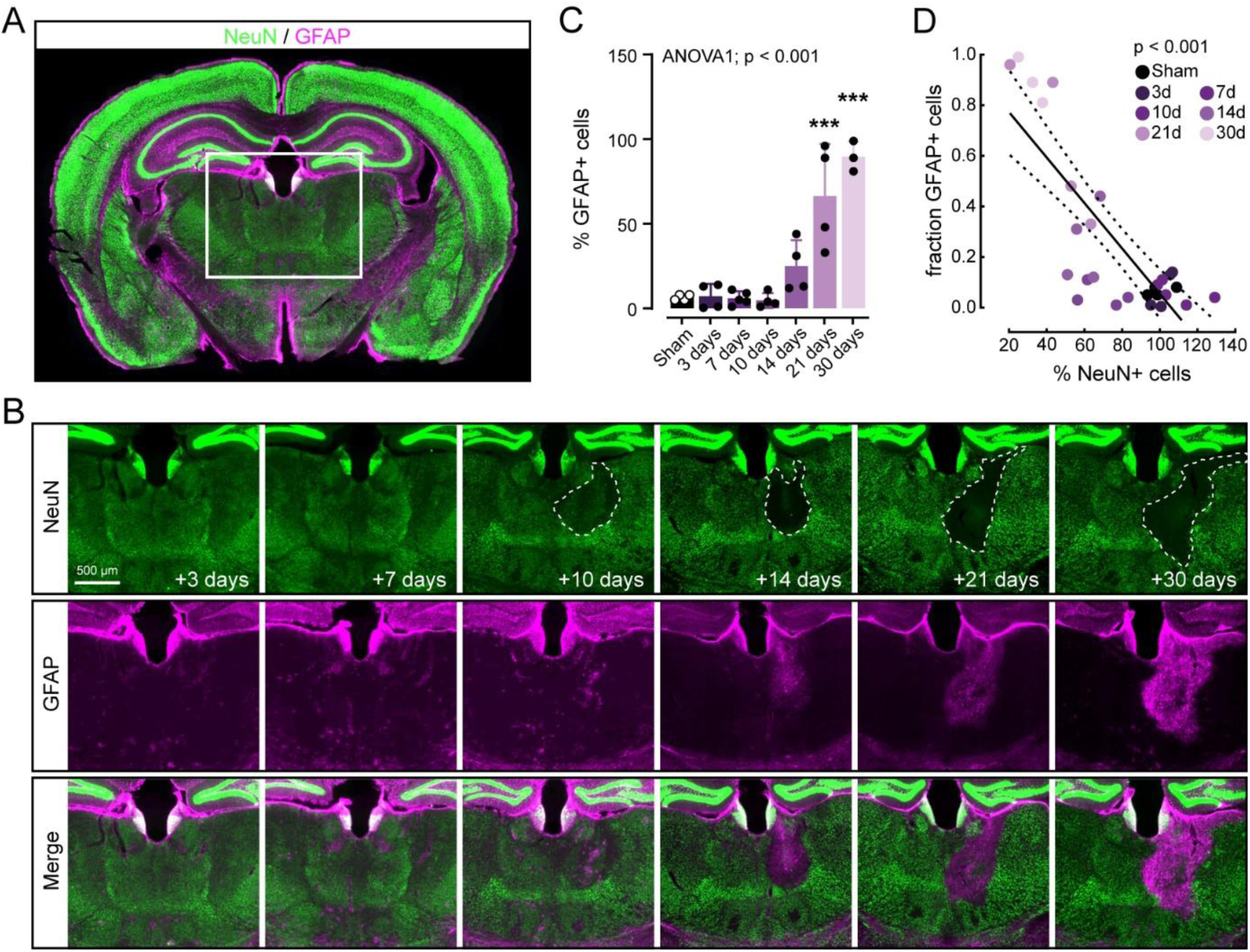
Cre-mediated glial scar. **A)** Representative epi-fluorescent image of a coronal brain slice taken from a mouse sacrificed 3 days after AAV1-CRE injection. Green: NeuN, purple: GFAP. **B)** Zoomed image of the thalamic region of coronal slices represented by the white square in A), of non-injected (left) and injected (right) MD at different timepoints post-injection of AAV1-CRE. Top: NeuN-immunostaining with a dashed line surrounding the area with a clear absence of NeuN-positive cells. Middle: GFAP-immunostaining showing an increase with time. Bottom: Merge of NeuN- and GFAP-immunostaining showing that the increase in GFAP happens where there are no NeuN-positive cells. **C)** % of GFAP-positive cells in the injected side of the MD and normalized by the number of DAPI-positive cells in the same region. Each dot corresponds to a mouse and error bars are sd. **D)** Correlation between GFAP-positive cells normalized to DAPI-positive cells in the MD at the different time points post-injection and the NeuN-positive cells normalized to DAPI-positive cells in the MD. Each dot corresponds to a mouse represented in C). Filled line is the best fit using a simple linear regression.

Next, we sought to confirm that the reduction in NeuN+ cells reflected true neuronal loss rather than mere downregulation of NeuN protein expression, and to determine whether similar patterns of neuronal loss / glial proliferation occurred following AAV1-CRE injections into other brain regions (**Fig.4**). To this end, AAV1-CRE was unilaterally injected into the ventrolateral (VL) thalamic nucleus of Thy1-GCaMP mice, which selectively express GCaMP in excitatory neurons^28,29^, and used GFP, a GCaMP core component, as an independent readout of neuronal survival (**Fig.4A, B**). As compared to DAPI+ cells, post-mortem GFP immunostaining revealed a marked and significant reduction in GFP+ cells in the injected VL, normalized to the non-injected contralateral side, which closely matched the decrease in NeuN+ cells (mean % ± sd; DAPI+: 107.2 ± 12.45; GFP: 46.32 ± 29.29; NeuN+ 46.04 ± 42.48; p=0.0017; one-way ANOVA) (**Fig.4C**). As in the MD, DAPI+ cell numbers remained stable between injected and non-injected VL (contralateral: 1090 ± 261.7sd; ipsilateral: 1155 ± 224.6; paired t-test). Together, these findings revealed a strong correlation between NeuN+ and GFP+ cells (Pearson’s correlation, p < 0.0001, r²=0.89), confirming that AAV1-CRE induces bona fide neuronal loss (**Fig.4E**).

**Figure 4.**
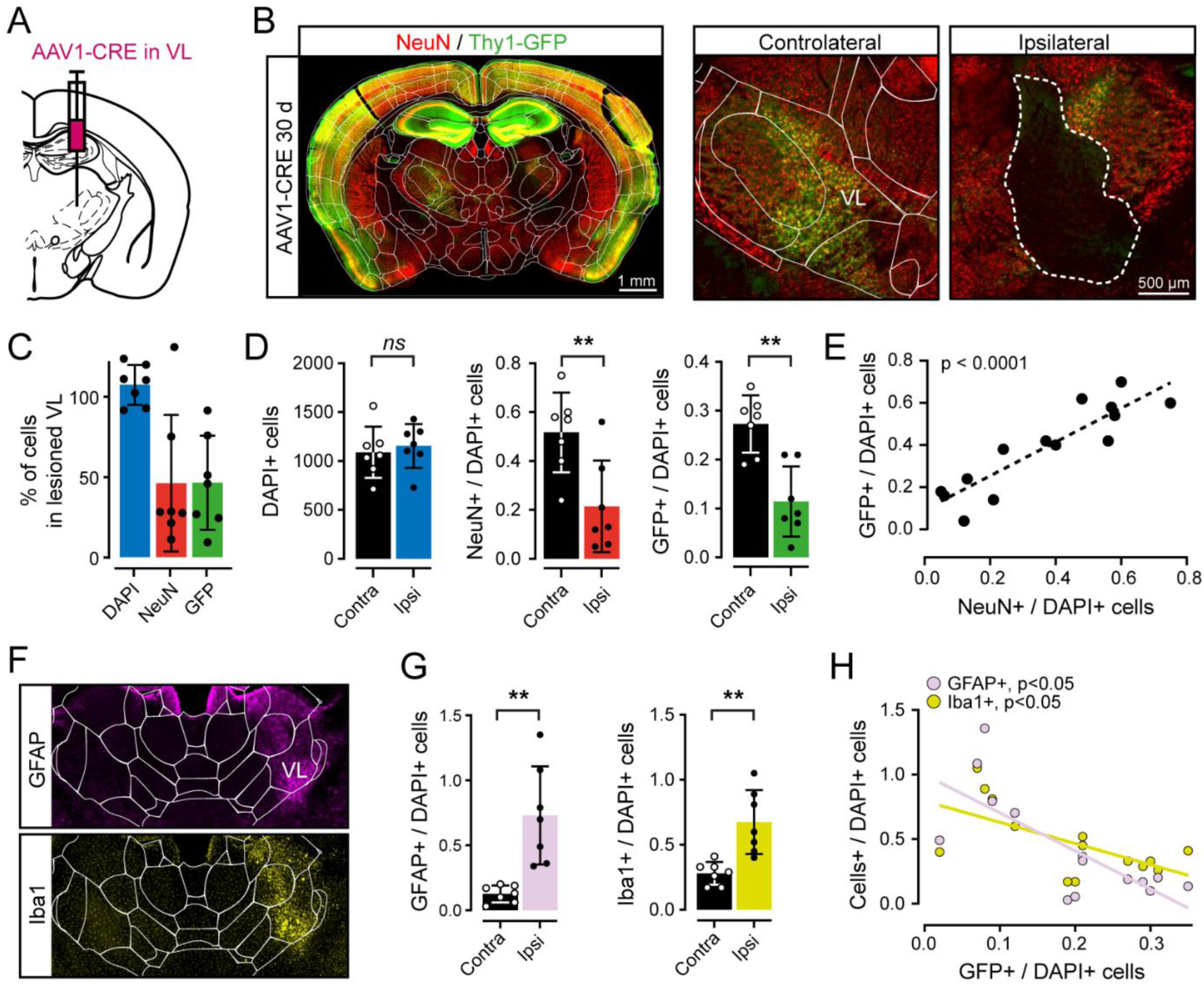
Cre-mediated loss of excitatory neurons. **A)** Schematic representation of the stereotaxic delivery of AAV1-CRE into the VL of Thy1 mice expressing GCaMP in excitatory neurons. **B)** Left: representative epi-fluorescent image of a coronal brain slice aligned to the Allen brain atlas taken from a mouse injected with AAV1-CRE into the VL of the right hemisphere, 30 days before sacrificed (30d). Red: NeuN, green: GFP. Right: zoomed image of the non-injected (contralateral) and injected (ipsilateral) VL from the left image, with a dashed line surrounding the area with a clear absent of GFP- and NeuN-positive cells. **C)** % of cells in the injected side normalized by the number of positive cells in the non-injected side of the VL sections measured by the Quint workflow (see methods) for the different fluorescent tags associated to the different somatic markers (DAPI, NeuN and GFP). Each dot corresponds to a mouse and error bars are sd. **D)** Left: number of DAPI-positive cells in the non-injected side (contra) and in the injected side (ipsi) of the VL sections with unilateral injection of AAV1-CRE in the VL. Middle: number of NeuN-positive cells normalized by the number of DAPI-positive cells in the non-injected side (contra) and in the injected side (ipsi) of the VL sections with unilateral injection of AAV1-CRE in the VL. Right: number of GFP-positive cells normalized by the number of DAPI-positive cells in the non-injected side (contra) and in the injected side (ipsi) of the VL sections with unilateral injection of AAV1-CRE in the VL. Each dot corresponds to a mouse and error bars are sd. **E)** Correlation between NeuN-positive cells normalized to DAPI-positive cells in the VL and the GFP-positive cells normalized to DAPI-positive cells in the VL. Each dot corresponds to a mouse represented in D). Dashed line is the best fit using a simple linear regression. **F)** Top: representative epi-fluorescent image of a coronal brain slice aligned to the Allen brain atlas, zoomed in the thalamic region and taken from a mouse injected with AAV1-CRE into the VL of the right hemisphere 30 days before, and with anti-GFAP immunostaining (purple). Bottom: representative epi-fluorescent image of the same brain slice represented on the Top, but with anti-Iba1 immunostaining (yellow). **G)** Left: number of GFAP-positive cells normalized by the number of DAPI-positive cells in the non-injected side (contra) and in the injected side (ipsi) of the VL sections with unilateral injection of AAV1-CRE in the VL. Right: number of Iba1-positive cells normalized by the number of DAPI-positive cells in the non-injected side (contra) and in the injected side (ipsi) of the VL sections with unilateral injection of AAV1-CRE in the VL. Each dot corresponds to a mouse and error bars are sd. **H)** Light pink: correlation between GFAP-positive cells normalized to DAPI-positive cells in the VL and the GFP-positive cells normalized to DAPI-positive cells in the VL. Yellow: correlation between Iba1-positive cells normalized to DAPI-positive cells in the VL and the GFP-positive cells normalized to DAPI-positive cells in the RTN. Each dot corresponds to a mouse represented in G). Lines represent the best fit using a simple linear regression.

To confirm the presence of astrocytes and microglia at the Cre-injected site, we quantified GFAP+ and Iba1+ cells (**Fig.4F**), which are established markers of astrocytes and microglia, respectively^26,27,30^. Similar to the MD, we observed a significant increase in GFAP+ cells within the injected VL compared to the non-injected side (mean GFAP+/DAPI+ cells ± sd; contralateral: 0.12 ±0.06; ipsilateral: 0.73 ± 0.38; paired t-test, p=0.002). This was accompanied by a comparable rise in Iba1+ cells (mean Iba1+/DAPI+ cells ± sd; contralateral: 0.28 ± 0.09; ipsilateral: 0.67 ± 0.24; paired t-test, p=0.003) (**Fig.4G**). Notably, the fraction of GFP+ neurons was significantly and inversely correlated with the fraction of both GFAP+ and Iba1+ cells (Pearson’s correlation; GFAP: p=0.0023, r²=0.55; Iba1: p=0.016, r^2^=0.39) (**Fig.4H**), indicating that neuronal loss is closely associated with the reactive proliferation of multiple glial cell types and occurs independently of the targeted brain area.

### Similar pattern of glial cell proliferation accompanies Cre-mediated loss of inhibitory neurons

Because the MD and VL thalamic nuclei are predominantly composed of excitatory neurons^31^ and GCaMP expression in Thy1-GCaMP mice is restricted to excitatory populations^28,29^, we next examined whether Cre recombinase also induces neurodegeneration within inhibitory neuronal populations (**Fig.5**). To this end, AAV1-CRE was unilaterally injected into the reticular thalamic nucleus (RTN) (**Fig.5A**). Unlike thalamic relay nuclei, the RTN is composed exclusively of inhibitory neurons, with no excitatory projection neurons. We assessed inhibitory neuronal survival by conducting post-mortem immunohistochemistry for NeuN and parvalbumin (PV), a marker of fast-spiking inhibitory neurons expressed by a substantial proportion of RTN neurons^32,33^ (**Fig.5B**). The fractions of PV+ and NeuN+ cells in the injected RTN relative to the non-injected control hemisphere were significantly smaller than the fraction of DAPI+ cells (mean % ± sd; DAPI: 92.8 ± 9.67; PV: 61.9 ± 9.05; NeuN: 62.75 ± 9.12; p<0.0001; one-way ANOVA) (**Fig.5C**). Similar to observations in the MD and VL regions, the number of DAPI+ cells did not differ significantly between Cre-injected and non-injected sites within the RTN (contralateral: 1959 ±241.5 ipsilateral: 1813 ± 254.7; paired t-test, p=0.12) (**Fig.5D**). Cre expression reduced the number of NeuN+ and PV+ cells relative to DAPI+ cell counts (**Fig.5D**), leading to an overall significant positive correlation between NeuN+ and PV+ cell numbers (Pearson’s correlation, p<0.0001, r² = 0.91) (**Fig.5E**). This finding indicates that Cre expression similarly affects both excitatory and inhibitory neurons, suggesting the involvement of shared cellular mechanisms. Consistent with this, we observed a significant increase in both GFAP+ and Iba1+ cells in the injected TRN (mean GFAP+/DAPI+ cells ± sd; contralateral: 0.32 ± 0.075; ipsilateral: 0.69 ± 0.09; paired t-test, p<0.0001; mean Iba1+/DAPI+ cells ± sd; contralateral: 0.23 ± 0.04; ipsilateral: 0.5 ± 0.11; paired t-test, p=0.0014) (**Fig.5F, G**). Furthermore, the increase in GFAP+ and Iba1+ cells was significantly and inversely correlated with the fraction of PV+ neurons (Pearson’s correlation; GFAP: p=0.0167, r²=0.39; Iba1: p=0.0147, r²=0.4) (**Fig.5H**), mirroring the pattern observed for putative excitatory neurons in the MD and VL nuclei.

**Figure 5.**
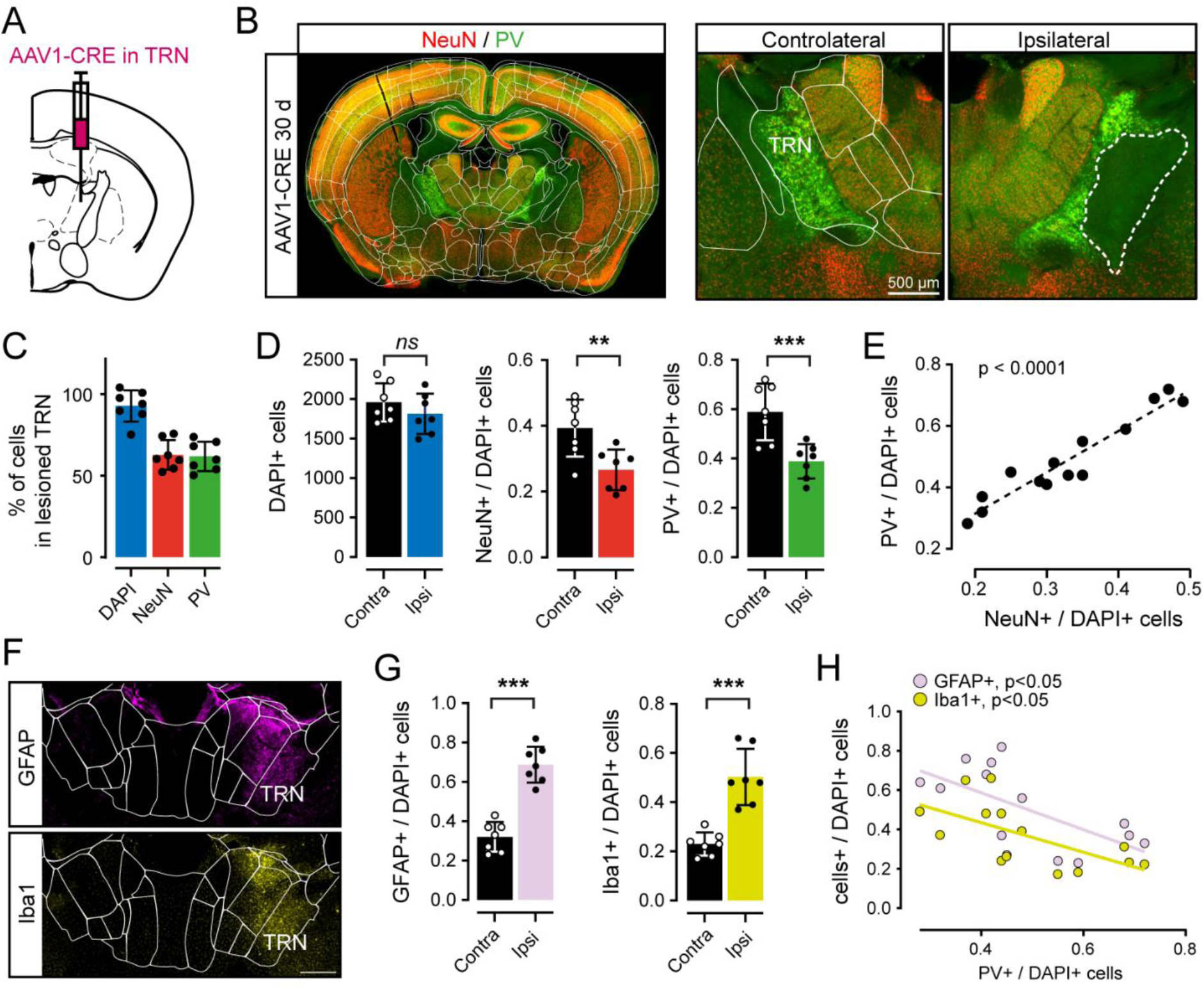
Cre-mediated loss of inhibitory neurons. **A)** Schematic representation of the stereotaxic delivery of AAV1-CRE into the RTN. **B)** Left: representative epi-fluorescent image of a coronal brain slice aligned to the Allen brain atlas taken from a mouse injected with AAV1-CRE into the RTN of the right hemisphere, 30 days before sacrificed (30d). Red: NeuN, green: PV. Right: zoomed image of the non-injected (contralateral) and injected (ipsilateral) RTN from the left image, with a dashed line surrounding the area with a clear absent of PV- and NeuN-positive cells. **C)** % of cells in the injected side normalized by the number of positive cells in the non-injected side of the RTN sections for the different fluorescent tags associated to the different somatic markers (DAPI, NeuN and PV). Each dot corresponds to a mouse and error bars are sd. **D)** Left: number of DAPI-positive cells in the non-injected side (contra) and in the injected side (ipsi) of the RTN sections with unilateral injection of AAV1-CRE in the RTN. Middle: number of NeuN-positive cells normalized by the number of DAPI-positive cells in the non-injected side (contra) and in the injected side (ipsi) of the RTN sections with unilateral injection of AAV1-CRE in the RTN. Right: number of PV-positive cells normalized by the number of DAPI-positive cells in the non-injected side (contra) and in the injected side (ipsi) of the RTN sections with unilateral injection of AAV1-CRE in the RTN. Each dot corresponds to a mouse and error bars are sd. **E)** Correlation between NeuN-positive cells normalized to DAPI-positive cells in the TRN and the PV-positive cells normalized to DAPI-positive cells in the RTN. Each dot corresponds to a mouse represented in D). Dashed line is the best fit using a simple linear regression. **F)** Top: representative epi-fluorescent image of a coronal brain slice aligned to the Allen brain atlas, zoomed in the thalamic region and taken from a mouse injected with AAV1-CRE into the RTN of the right hemisphere 30 days before, and with anti-GFAP immunostaining (purple). Bottom: representative epi-fluorescent image of the same brain slice represented on the Top, but with anti-Iba1 immunostaining (yellow). **G)** Left: number of GFAP-positive cells normalized by the number of DAPI-positive cells in the non-injected side (contra) and in the injected side (ipsi) of the RTN sections with unilateral injection of AAV1-CRE in the RTN. Right: number of Iba1-positive cells normalized by the number of DAPI-positive cells in the non-injected side (contra) and in the injected side (ipsi) of the RTN sections with unilateral injection of AAV1-CRE in the RTN. Each dot corresponds to a mouse and error bars are sd. **H)** Light pink: correlation between GFAP-positive cells normalized to DAPI-positive cells in the TRN and the PV-positive cells normalized to DAPI-positive cells in the RTN. Yellow: correlation between Iba1-positive cells normalized to DAPI-positive cells in the RTN and the PV-positive cells normalized to DAPI-positive cells in the RTN. Each dot corresponds to a mouse represented in G). Lines represent the best fit using a simple linear regression.

### Cre-mediated neuronal loss impairs associative learning independent of viral serotype

To investigate the roles of viral serotype and Cre expression in neuronal loss and associated phenotypic changes, we unilaterally injected AAV vectors encoding Cre recombinase into the basolateral amygdala (BLA) of C57BL/6 mice, using two different serotypes: AAV1 and AAV5, which differ in their tissue tropism and spread within the brain^34,35^. To control for potential effects of a fluorescent reporter, we used an AAV5 vector encoding GFP. As a control condition, we also injected AAV1-GCaMP7f, a widely used vector for calcium imaging that does not induce Cre-related toxicity^36,37^ (**Fig.6A**). Post-mortem NeuN immunostaining revealed a marked and highly significant reduction in the number of NeuN+ cells in AAV1-CRE-injected mice compared to AAV1-GCaMP7f-injected controls (mean fraction of NeuN+ positive cells ± sd; AAV1-GCaMP7f: 1.03± 0.11; AAV1-CRE: 0.77 ± 0.15; p=0.0001) (**Fig.6B, C**). In contrast, AAV5-CRE-injected mice exhibited a less significant reduction in NeuN+ cell numbers compared to controls (AAV1-GCaMP7f: 1.03± 0.11; AAV5-CRE: 0.89 ± 0.041; p=0.04) (**Fig.6C**), likely reflecting the lower transduction efficiency and more restricted spread of AAV5 relative to AAV1^34,35^, resulting in a smaller but still significant lesion (**Fig.6B**).

**Figure 6.**
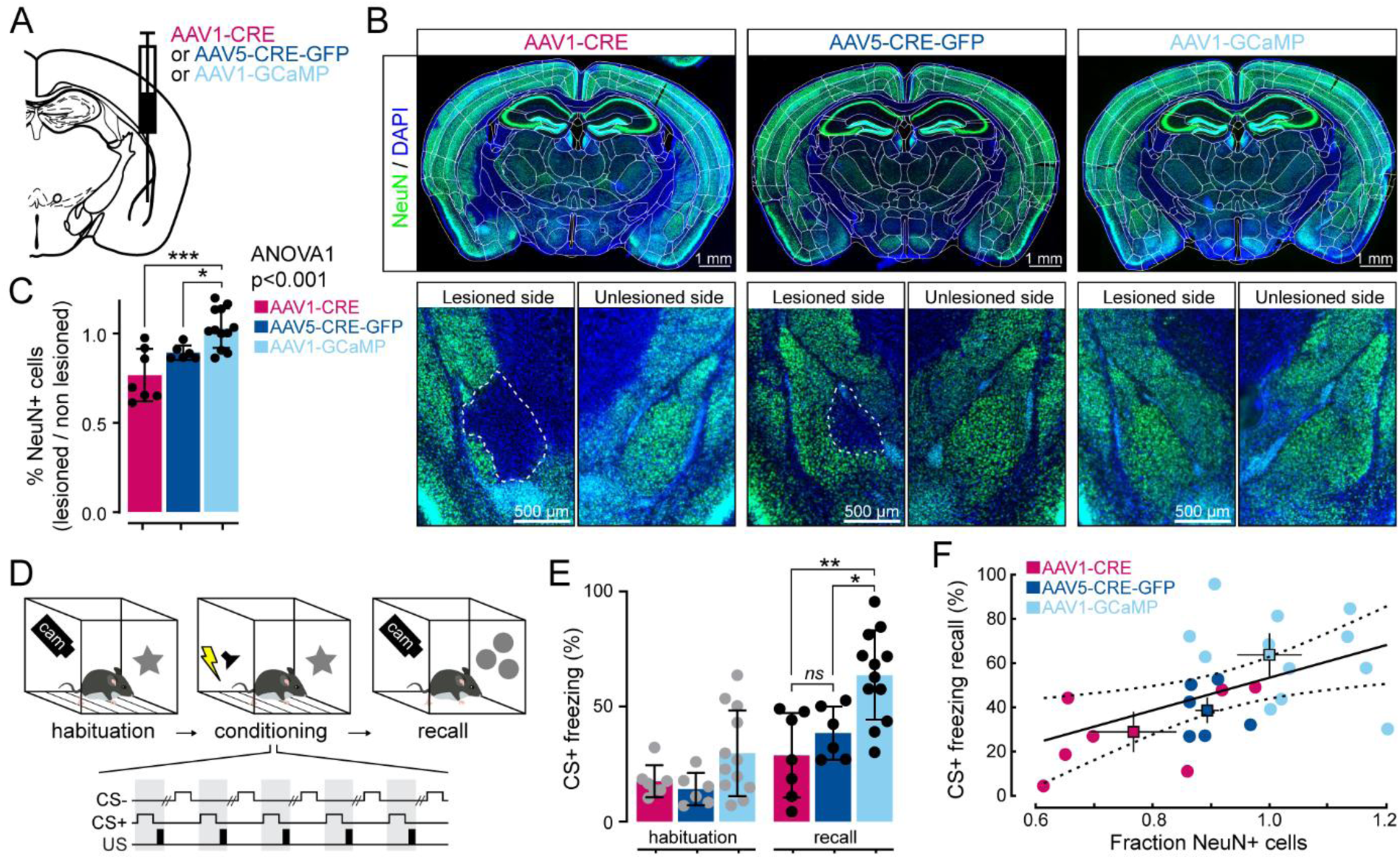
Cre-mediated neuronal loss impairs associative learning. **A)** Schematic representation of the stereotaxic delivery of different AAVs into the BLA. **B)** Top: tree epifluorescent images of coronal brain slices aligned to the Allen brain atlas and taken from mice injected with various AAV serotypes as indicated in the panels. Blue: DAPI, green: NeuN. Bottom: zoomed images of the lesioned and non-lesioned BLA from the top images, respectively, with a dashed line surrounding the area with a clear absent of NeuN-positive cells. **C)** % of NeuN-positive cells in the lesioned side normalized by the number of NeuN-positive cells in the non-lesioned side for BLA sections for the different AAVs injected into the BLA. Each dot corresponds to a mouse and error bars are sd. **D)** Schematic representation of the auditory fear conditioning protocol used to test associative learning. **E)** % of freezing during the first 2 CS+ presentation during habituation and recall in the different groups of mice injected with the different AAVs represented in A). Each dot corresponds to a mouse and error bars are sd. **F)** Correlation between fraction of NeuN-positive cells in the BLA represented in C) and % of freezing during the CS+ of the recall represented in E). Each dot corresponds to a mouse, squares represent the mean of each group and error bars are sd. Filled line is the best fit using a simple linear regression.

We next investigated whether BLA neuronal loss induced by AAV1-CRE or AAV5-CRE leads to detectable behavioral alterations. Given the critical role of BLA neurons in learned fear-related behaviors^38–40^, we performed auditory fear conditioning 21 days after AAV-CRE injection with both serotypes (**Fig.6D**), prior to NeuN immunostaining. Mice injected with AAV1-GCaMP7f in the BLA served as controls, as no lesions were observed in this group (**Fig.6B, C**). Auditory fear conditioning was performed using a classical discriminative protocol, as previously described^40^. In brief, mice were exposed to five presentations of auditory stimuli: an 8 kHz tone (CS+) paired with a mild foot shock (0.6 mA) and a white Gaussian noise stimulus (CS-) presented without shock, delivered in a pseudorandom sequence (**Fig.6D, Methods**). To evaluate fear learning, freezing responses to the CS+ were measured during a recall session conducted 24 hours after conditioning (**Fig.6D**).

During the habituation phase, animals across all groups exhibited comparable levels of CS+-evoked freezing (mean ± sd: AAV1-GCaMP7f, 29.76 ± 18.62%; AAV1-CRE, 17.61 ± 7.0%; AAV5-CRE, 14.30 ± 7.08%; one-way ANOVA, p=0.0687) (**Fig.6E**), indicating no baseline differences in fear responses prior to conditioning. However, these responses changed markedly during the recall session, with freezing behavior that differed significantly between groups. Animals injected with AAV1-GCaMP7f showed significantly higher freezing levels (mean ± sd: AAV1-GCaMP7f: 63.7 ± 19.4%; AAV1-CRE: 28.8 ± 18.4%; AAV5-CRE: 38.6 ± 11.5%; one-way ANOVA, p<0.0001) (**Fig.6E**). These results confirm that AAV1-GCaMP7f expression did not disrupt BLA neuronal circuits or impair fear learning behavior. Interestingly, although the lesions induced by AAV5-CRE were smaller than those caused by AAV1-CRE (**Fig.6B, C**), freezing levels in AAV5-CRE-injected mice were still significantly lower than controls (mean ± sd: AAV1-GCaMP7f: 63.75 ± 19.40% vs. AAV5-CRE: 38.57 ± 11.50%; p=0.0321) and closely matched the reduced freezing seen in AAV1-CRE mice (mean ± sd: AAV1-CRE: 28.85 ± 18.38%; p=0.88) (**Fig.6E**). These results suggest that the impairment in fear learning is not driven by viral serotype or tropism, but by Cre expression itself. Supporting this conclusion, we found a significant positive correlation between the fraction of surviving NeuN+ neurons and freezing behavior during recall (Pearson’s correlation, p=0.0115, r²=0.247), indicating that greater neuronal loss was associated with more pronounced behavioral deficits, independent of the viral serotype used. (**Fig.6F**). Taken together, these findings demonstrate that it is the expression of Cre, rather than AAV infection or associated fluorescent tags, that drives local neuronal degeneration and leads to impaired fear learning.

## DISCUSSION

In this study, we demonstrate that Cre recombinase expression alone, in the absence of LoxP sequences, is sufficient to induce a reproducible focal lesion in the central nervous system. The mechanisms underlying Cre-induced neurodegeneration are not fully understood, but evidence points to DNA damage as a central factor. Cre recombinase is known to induce double-strand breaks at cryptic or pseudo-LoxP sites distributed throughout the genome^41^, leading to genomic instability, cell cycle arrest, and activation of apoptotic pathways. This off-target genotoxic effect appears to be dose-dependent^16,20^, but with high titration, occurs regardless of the AAV serotype used and does not require a preceding neuroinflammatory response to initiate neurodegeneration. Notably, Cre-induced neuronal loss impacts both excitatory and inhibitory populations and spans multiple brain regions, including the mediodorsal thalamus (MD), ventral lateral thalamic nuclei (VL), reticular thalamic nucleus (RTN), and basolateral amygdala (BLA). These results align with recent findings reporting Cre toxicity in thalamic^18^ and dopaminergic neurons^16,17^, highlighting a broad vulnerability of CNS neurons to high levels of Cre expression.

The lesion induced by Cre expression develops progressively, with detectable neuronal loss emerging around 10 days post-injection and increasing thereafter. Notably, these focal lesions are not apparent with standard DAPI staining and only become evident when using neuron-specific markers such as NeuN or PV. This highlights the critical importance of employing cell-type-specific markers when evaluating the effects of intracerebral manipulations, as relying on general nuclear stains may overlook subtle, yet biologically significant changes. Importantly, Cre expression is detectable as early as 3 days post-injection, preceding any visible neuronal loss. This is evidenced not only by direct detection of the Cre protein but also by the robust increase in TdTomato fluorescence in transgenic Ai9 mice harboring *loxP* sites. Our findings indicate that neurodegeneration begins only after Cre expression reaches a high threshold, peaking around 10 days post-injection, consistent with previous reports linking Cre toxicity to elevated expression levels^20,42^. These results suggest two possibilities: (1) neurons may tolerate low to moderate levels of Cre expression without immediate degeneration, and/or (2) Cre-mediated neuronal loss requires several days to manifest after reaching cytotoxic levels. Either way, our data define a critical time window between 3 and 10 days post-injection during which Cre is actively expressed, but significant neuronal loss has not yet occurred. This window may offer a valuable opportunity to assess Cre-mediated effects, while minimizing confounding impacts of Cre toxicity.

Nevertheless, we observed a lack of correlation between Cre expression levels and the level of its downstream reporter TdTomato, suggesting that reporter signal alone is an unreliable indicator of viable Cre-expressing neurons. Instead, TdTomato fluorescence appears to persist or even accumulate in lesioned tissue with undergoing neuronal degeneration. This supports a model in which Cre-induced DNA damage leads to apoptosis of neurons^16,41^, followed by clearance through glial phagocytosis. Supporting this, we observed TdTomato signal in cells with glial morphology, alongside increased Iba1 and GFAP levels following the establishment of the focal lesion. This suggests that microglia and astrocytes are recruited to the site of degeneration and may internalize neuronal debris, including fluorescent reporter proteins, during phagocytosis. Such transfer of fluorescent signal to glial cells has been reported in models of neuronal injury and tauopathy, where microglia engulf labeled neurons and retain their fluorescence^43,44^. These findings underscore the importance of distinguishing between true transgene expression and signal persistence due to phagocytic uptake, specifically when interpreting cell-type specificity based solely on Cre reporter labeling.

The recruitment of astrocytes and microglia occurs subsequent to the establishment of the Cre-induced focal lesion, rather than preceding it. Notably, we observed that classical neuroinflammation markers such as GFAP and Iba1 only increased after significant neuronal loss had occurred. This temporal pattern indicates that the neuroinflammatory response is a consequence and not a cause of neurodegeneration in this model. These findings contrast with other neurodegenerative contexts where inflammation can precede and even contribute to neuronal loss^45,46^. Supporting this, we found that viral delivery of GCaMP using the same AAV vector backbone did not lead to neuronal death, underscoring that the observed neurodegeneration is specific to Cre expression rather than a nonspecific response to viral infection. This is consistent with prior studies showing that AAV vectors themselves are generally well tolerated and do not induce innate immune responses or cytotoxicity at moderate titers^47,48^. Importantly, the delayed upregulation of GFAP and Iba1 is characteristic of a glial response to tissue damage and is consistent with reactive gliosis and scar formation following CNS injury^25^. These findings suggest that the astrocytic and microglial activation observed in our model reflects a secondary clearance response to Cre-induced neuronal death, rather than an initiating factor.

In summary, our findings demonstrate that viral delivery of Cre recombinase provides an effective and controllable strategy for inducing focal lesions in the central nervous system. By selecting appropriate promoters to drive Cre expression, this approach allows for the targeted ablation of specific neuronal populations. Moreover, lesion size and anatomical precision can be finely tuned through adjustments in viral titer and injection volume, enabling localized manipulation of discrete brain regions, such as individual subcortical nuclei or cortical layers. This strategy offers a valuable tool for investigating the behavioral and functional consequences of region- or cell type-specific lesions. For example, we observed impairments in fear learning following neuronal loss in the BLA, while motor dysfunction was associated with Cre-mediated dopaminergic neuron loss in the ventral tegmental area^16,17^. Importantly, Cre-induced lesions present a more refined and ethically favorable alternative to traditional chemical lesion methods, such as those involving excitotoxins. Unlike excitotoxic lesions, Cre expression does not provoke seizures or require extensive post-procedure recovery, reducing animal distress and improving experimental reproducibility. Taken together, this approach represents a versatile and ethically sound method for modeling focal CNS injury and studying the underlying neural substrates of behavior.

## RESOURCES AVAILABILITY

Correspondence and reasonable requests for materials and data supporting this study should be addressed to. All data are available in the figures. Any additional information required to reanalyze the data reported in this work is available from the corresponding authors contact upon request.

## Acknowledgements

We thank H. El Oussini, P. Costet and C. Martin (PIV-EOPS & PIV-EXPE, Univ. Bordeaux, CNRS) for support with animal husbandry; the Bordeaux Imaging Center (BIC, Univ. Bordeaux, CNRS, INSERM, BIC, US4, UAR 3420), and all the members of the Gambino laboratory for assistance and helpful discussions.

FG received funding from: the European Research Council (ERC) under the European Union’s HORIZON research and innovation program (ERC-CoG-2021; MOTORHEAD; grant agreement 101043602); the French National Research Agency (ANR-21-CE37-0010-SyTUNE; ANR-22-CE37-0015-THALACOR; ANR-22-ERCC-0010-MOTORHEAD); the University of Bordeaux; and the Region Nouvelle Aquitaine. EA is supported by a Marie Skłodowska-Curie individual fellowship (HORIZON-MSCA-2023-PF-01 DANAL 101155516). This work was supported by the Bordeaux Neurocampus core facilities (LabEx BRAIN; ANR-10-LABX-43).

## AUTHOR CONTRIBUTIONS

Conceptualization, MG, EA, and FG. Methodology, MG, EA and FG. Investigation, MG, EA, and EC. Formal analysis, MG, EA, and FG. Resources, FG. Writing – original draft, review & editing, EA and FG. Visualization, MG, EA, and FG. Supervision, EA and FG. Funding acquisition, EA and FG. Project administration, FG.

## METHODS

### Experimental Model and Study Participant Details

A total of 16 males and 10 females Ai9Tomato for MD lesions, 3 males and 4 females Thy1GCaMP for VL and TRN lesions, 14 males and 14 females C57BL/6J from Charles River for BLA lesions, 2-7 months old, were used in this study. Mice were housed with littermates (2–4 mice per cage) in ventilated cages under an inverted 12 h dark/12 h light cycle, temperature was maintained between 19 °C and 23 °C and humidity between 50% and 65%. The mice were housed in enriched cages and provided with food and water ad libitum.

All experimental procedures were approved and performed in accordance with the local ethics and welfare Committees guidelines (N °50DIR_15-A) and by the French Ministry for Agriculture/Research (APAFIS #21728), in agreement with the European Communities Council Directive of September 22th 2010595(2010/63/EU, 74).

### Surgery and virus strain

All surgeries used standard aseptic procedures. Mice were deeply anesthetized with 4% of isoflurane (by volume in O2), or with an intraperitoneal injection of a mix containing medetomidine (sededorm,0.27 mg.kg-1) and buprenorphine (0.05 mg.kg-1) in sterile NaCl 0.9%, and mounted in a stereotaxic apparatus (RWD). Mice were kept on a heating pad and their eyes were covered with eye ointment ophthalmic gel. The scalp was shaved and disinfected with 70% ethanol and betadine. Local analgesia was achieved by application of 100 μl of lidocaïne (lurocaïne, 1%) injected subcutaneously in the scalp. 40 μL of dexamethasone (dexadreson, non-steroidal anti-inflammatory and analgesic drug, 5 mg.kg−1) was administrated intramuscularly in the quadriceps to prevent inflammation. A 1 cm incision in the skin was done and the skull was cleaned and dried with sterile cotton swabs. For stereotaxic injections the bregma and lambda were aligned (x and z). The injections targeted the MD (1 injection site: -1.0 mm AP; -3.4 mm DV; -0.4 mm ML from bregma), the VL (1 injection site: -0.45 mm AP; -3.75 mm DV; -1.3 mm ML from bregma), the RTN (1 injection site: -0.3 mm AP; -3.5 mm DV; -1.3 mm ML from bregma) or the BLA (2 injection sites: -1.44 mm AP; - 4.85 mm and -4.50 mm DV; - 2.89 mm ML from bregma). A total of 200nL (100 nL per site) of virus was injected at a maximum rate of 50 nL.min−1, using a glasspipette (Wiretrol, Drummond) attached to an oil hydraulic manipulator (MO-10, Narishige). AAV1.Syn.GCaMP7f.WPRE ([1e+13] titration), AAV1.hSyn.CRE.WPRE ([5e+13]) or AAV5.hSyn.CRE.eGFP ([1.7e+13] titration) were injected, all from Addgene, USA. After injections, the viruses were allowed to diffuse for 10 min before the pipette was withdrawn. At the end of the surgery, anesthesia was reversed by a sub-cutaneous injection of a mixture containing atipamezole (revertor, 2.5 mg.kg-1) and buprenorphine (buprecare, 0.1 mg.kg-1) in sterile 0.9% NaCl or by stopping the isoflurane.

### Histology and imaging

To evaluate the viral expression and lesion profiles, mice were deeply anesthetized with a mix of ketamine (100 mg.Kg-1) and xylazine (20 mg.Kg-1) and perfused with 4% paraformaldehyde. The brain was extracted and fixed for 24 h in paraformaldehyde at 4 °C, and then washed with phosphate-buffered saline.

The brain was sectioned at 60 μm, and immunohistology was done with primary antibodies: rabbit anti-NeuN (1:1000, Synaptic System), chicken anti-GFP (1:1000, AbCam), rabbit anti-CRE (1:2000, AbCam), chicken anti-GFAP (1:1000, AbCam), rabbit anti-Iba1 (1:1000, FUJIFILM Wako Pure Chemical Corporation), chicken anti-parvalbumine (1:2000, Novus Biologicals), and secondary antibodies: goat anti-rabbit 488 (1:500, Thermo Fisher Scientific), goat anti-chicken 647 (1:500, Thermo Fisher Scientific), goat anti-chicken 594 (1:500, Thermo Fisher Scientific). Briefly, free floating slices were washed and permeabilized in phosphate buffer solution with 0.1% triton X-100 (PBS-T), 3 times for 10 min each, at RT and under agitation. Further permeabilization and non-specific sites were blocked by incubating the slices for 1h at RT under agitation in blocking solution (phosphate buffer solution with 5% normal goat serum (Grosseron or AbCam) and 0.3% triton X-100). Slices were then incubated for 12-14h at 4°C and under agitation with primary antibodies diluted in blocking solution. Afterwards, the slices were washed 3 times in PBS-T and incubated with the secondary antibodies diluted in blocking solution, for 2h at room temperature under agitation. Slices where then washed twice in PBS-T and once in PBS and mounted on glass slides and stained with DAPI.

Images were taken at ×20 magnifications for each section using a Hamamatsu Nanozoomer at different excitation wavelengths (405 nm, 488 nm, 570 nm and 647 nm for DAPI, FITC, TRITC and Cy5 respectively).

### Quantification

To register optical sliced to the Allen Brain Institute Taxonomy and quantify all different immunostainings throughout the different experiments (NeuN, DAPI, Cre, tdTomato expression, GFAP, Iba1, GFP, PV), we used the DeepSlice software as well as the QuintWorkflow. In quick, DeepSlice was used to automatically register mouse brain histological images to the Allen Brain Common Coordinate Framework. The results were then passed on the QuickNII software to grossly adjust the slices to the Allen Brain atlas, and then more detailly by using the VisuAlign software. The Ilastic software was trained to segment cell and background for all the different immunostainings. The QNLMask software was used to separate the hemispheres on the maps obtained through VisuAlign, allowing us to do cell quantification on each side of the slices. Finally, the Nutil software regroups all the previous results in order to give an output in which for each brain region there is the quantification for each cell type.

### Behavior

At least 5 days before starting behavioral experiments, mice went through handling with the same experimenter that performed the experiments in order to decrease stress. For consistency across experiments, mice were then habituated to auditory tones. During habituation, mice were placed on the conditioning compartment (context A, consisting of a squared box with a grid floor that allows the delivery of a foot shock and with home cage litter under; cleaned between individuals with 70% ethanol). Two conditional auditory stimuli (CS) (8 kHz pure tone; and white Gaussian noise (WGN); each composed of 27 pips, 50 ms in duration, 0.9 Hz for 30 s) were presented 4 times each with an 80-dB sound pressure level and variable inter stimulus interval (ISI). The freezing time during each CS presentation was measured and the mice returned to their home cage. Mice were fear conditioned 24 hours after the last habituation phase by using a classical differential protocol. Briefly, mice were exposed to context A and 5 auditory tones (CS+) were paired with the unconditional stimulus (US, 1s foot-shock, 0.6 mA). The onset of US coincided with the CS+ offset. 5 CS-presentations were intermingled with CS+ presentations with a variable (10-60 s) ISI. 8 kHz pure tones was used as CS+ and WGN was used as CS-. Recall tests were carried out 24 hours after the conditioning phase by measuring the freezing time during the presentation of 2 CS+ and 2 CS- in a new context (context B, consisting of a cylindrical white compartment with home cage litter on the floor; cleaned between individuals). For each behavioral session, the total time duration (sec) of freezing episodes upon CS+ and CS-presentation was quantified automatically using a fire-wire CCD-camera connected to an automated freezing detection software (POLY, Imetronic), and expressed as % of freezing.

### Statistics

Results are presented as mean ± standard deviation (sd) as indicated in the corresponding results section. Statistical analyses were performed using the software GraphPadPrism. To test for differences between several groups (figures 1, 2, 3 and 6), we used one-way ANOVA followed by a Dunnett’s multiple comparison test using sham as control for figures 1-3 and AAV1.GCaMP-injected mice in figure 6. To compare between injected and non-injected sides (figures 4 and 5) we used paired t-test. In all figures, correlation was tested using Pearson’s test.

